# Frequency-and circuit-specific effects of septohippocampal deep brain stimulation in mice as measured by functional ultrasound imaging

**DOI:** 10.1101/2023.05.21.541598

**Authors:** Lindsey M. Crown, Kofi Agyeman, Wooseong Choi, Nancy Zepeda, Ege Iseri, Pooyan Pahlavan, Steven J. Siegel, Charles Liu, Vasileios Christopoulos, Darrin J. Lee

## Abstract

**Background:** Deep brain stimulation (DBS) has shown remarkable success in treating neurological and psychiatric disorders such as Parkinson’s disease, dystonia, epilepsy, and obsessive-compulsive disorder. Despite this success, the underlying mechanism of action remains unknown. DBS is now being explored to improve functional outcomes in other psychiatric conditions, such as those characterized by reduced N-methyl-D-aspartate (NMDA) function (i.e. schizophrenia). While DBS for movement disorders requires high-frequency continuous stimulation, there is evidence that intermittent low-frequency stimulation in neuropsychiatric conditions may have persisting cognitive benefits, necessitating a broader exploration of how DBS alters brain networks.

**Objective:** We characterize the effects of pharmacologic NMDA antagonism on the septohippocampal network and the impact of high- and low-frequency MSN DBS on cerebral blood volume (CBV) in brain structures within and outside of the septohippocampal network.

**Methods:** In this study, we utilize a novel technology, functional ultrasound imaging (fUSI), to characterize the cerebrovascular impact of medial septal nucleus (MSN) DBS under conditions of NMDA antagonism (pharmacologically using Dizocilpine [MK-801]) in anesthetized male mice.

**Results:** Imaging from a sagittal plane across a variety of brain regions, we find that MSN theta-frequency (7.7Hz) DBS has a larger effect on hippocampal CBV after stimulation offset. This is observed following an intraperitoneal (i.p.) injection of either saline vehicle or MK-801 (1 mg/kg). This effect is not present using standard high-frequency DBS stimulation parameters (i.e. gamma [100Hz]).

**Conclusion:** These results indicate the MSN DBS increases circuit-specific hippocampal neurovascular activity in a frequency-dependent manner that continues beyond the period of electrical stimulation.

## Introduction

There is growing interest in utilizing neuromodulation to treat cognitive impairment associated with neurological and psychiatric disorders. Many of these disorders involve aberrant electrophysiology and cerebral blood perfusion [1,2]. Recent evidence demonstrates that electrical neuromodulation can improve functional outcomes [3,4]. Particularly, deep brain stimulation (DBS) has shown remarkable success in treating neurological diseases such as movement disorders and epilepsy, and there is increasing evidence for its efficacy in cognitive and psychiatric conditions.

Despite this growing utility, the underlying mechanism of DBS for cognitive outcomes remains largely unknown. Pre-clinical studies have been hindered by technological limitations, such as the inability to record electrical brain activity during stimulation and the low spatial resolution of electrographic measures. Functional ultrasound imaging (fUSI) is a relatively new technology that enables large-scale estimates of neural activity through measures of cerebral blood volume (CBV). fUSI provides a unique combination of high spatiotemporal resolution (∼100 μm^3^, up to 10 ms) and high sensitivity to slow blood flow (∼1 mm/s velocity) across a large field of view. In fact, fUSI has already been proven to be an effective tool for imaging large-scale brain activity and pharmacodynamics [5–8]. As such, it is well-positioned to improve our understanding of the impact of DBS on large-scale brain networks during and immediately following stimulation.

In many disorders, cognitive dysfunction is accompanied by altered septohippocampal network activity. For instance, neural oscillatory patterns within the septohippocampal network, such as gamma- and theta-band activity, are characteristically altered in disorders such as schizophrenia and Alzheimer’s disease and are often correlated with impaired memory [1,9–11]. Alterations in specific neurotransmitter signal transduction pathways also play an important role in modulating neural oscillatory activity. In particular, the glutamatergic *N*-methyl-D-aspartate (NMDA) receptor is crucial to regulating hippocampal theta and gamma oscillations and is the predominant molecular control for synaptic plasticity and memory function [12–14] As such, pharmacologic NMDA receptor antagonism (i.e. via MK-801) results in characteristic changes to neural oscillatory patterns and memory dysfunction [15–19].

The medial septal nucleus (MSN) is a key structure in the septohippocampal network that modulates sensory-motor processing and acts as a “pacemaker” for hippocampal theta oscillations via dense glutamatergic, cholinergic, and GABAergic projections to the hippocampus [20,21]. This makes the MSN a promising target for DBS in cognitive disorders involving memory impairments [22]. Our recent work as well as that of others suggest that modulating the septohippocampal network via MSN theta frequency-specific (7.7 Hz) DBS can restore cognitive impairment and memory dysfunction in preclinical models of epilepsy, traumatic brain injury, Alzheimer’s disease, and schizophrenia [22–27].

In the current study, we utilize fUSI to characterize the effects of MK-801 on CBV in the septohippocampal network (Fig. 1A) including the hippocampus, and medial prefrontal cortex (mPFC). Within the same sagittal plane, we also image CBV changes (ΔCBV) to regions of interest (ROIs) outside this network including the striatum, thalamus, hypothalamus, and pallidum. Note that other regions that are connected with the MSN, such as amygdala, habenula, or raphe nucleus were not recorded, since they were not accessible from the selected 2D image plane. Additionally, we determine the effect of direct MSN stimulation on blood flow in these areas using two distinct frequencies, theta (7.7Hz) and gamma (100Hz). Finally, we test the hypothesis that MSN theta-frequency-specific stimulation can improve blood flow under conditions of reduced NMDA function.

**Figure 1.**
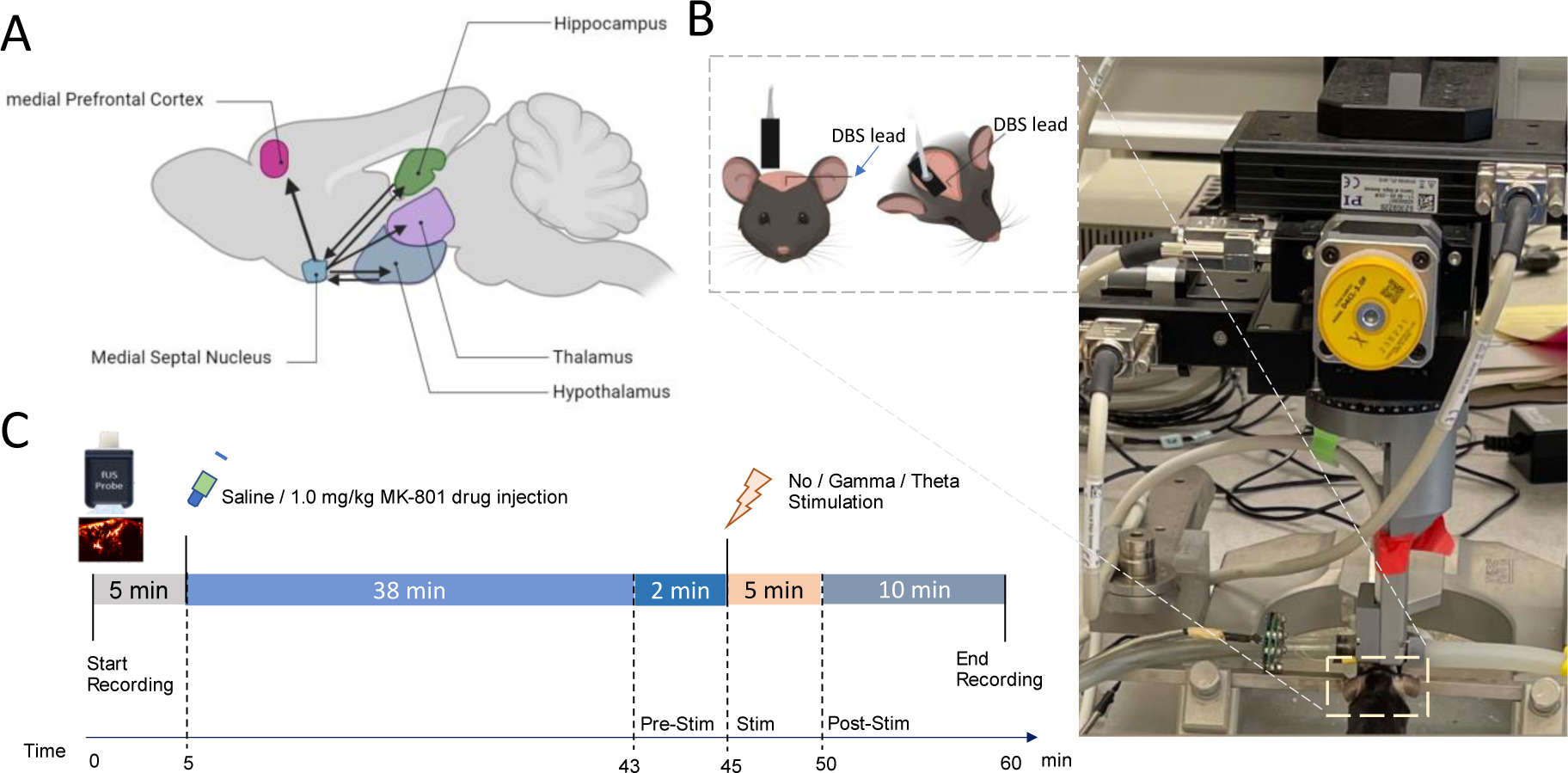
Experimental setup and fUSI recording protocol. **A)** Schematic illustration of connectivity between the MSN and ROIs. Arrowheads represent axonal projections to and/or from MSN. **B)** Experimental set-up showing the anesthetized mouse in a stereotaxic frame under the Iconeous One motorized probe mount. DBS stimulating electrodes were implanted on the left hemisphere and a sagittal plane of the right hemisphere was imaged**. C)** Diagram of the protocol for 60 minutes of continuous fUSI acquisition consisting of saline or 1.0 mg/kg MK-801 drug injection (at the 5-minute mark) and 5 minutes of gamma- or theta-frequency DBS (at the 45-minute mark).

## Materials and Methods

### Animals and surgical procedures

82 male 8–12-week-old C57BL/6 mice (Charles River Laboratories; Hollister, CA) were used in this study. fUSI data from two animals were excluded due to extreme values (Grubbs test for outliers, 98^th^ percentile of all maximum change in pD intensities) that did not appear physiological [28,29]. The animals were divided into two main groups: saline vehicle control (n=46) and MK-801 drug-administered (n=34). Each group was then sub-divided into three categories: no stimulation (saline: n=14; MK-801: n=10), theta stimulation (saline: n=16, MK-801: n=12), and gamma stimulation (saline: n=14, MK-801: n=14).

Mice were anaesthetized with 5% isoflurane in O_2_/N_2_O (1:2) carrier gas and then maintained at a constant rate (1.5-2%) through surgery and data acquisition. Body temperature was kept constant throughout recordings by placing animals on an electric heating pad. Hair was removed from the mouse’s head for fUSI using a commercially available depilatory cream (Nair, Pharmapacks).

To implant DBS electrodes, mice were head-fixed in a stereotaxic frame (David Kopf instruments, Tujunga, CA) and a midline incision of the scalp was made to expose the skull. A 2mm burr hole was then drilled to implant bipolar stimulating electrodes (E363T/2/SPC ELEC .008”/.2MM, Plastics One Inc., Roanoke, VA) targeting the midline MSN (AP: +0.7mm, ML: - 0.9mm, from bregma. Z: -4.39mm at 11.8 degrees) from the left hemisphere. Prior to implantation, the electrodes were connected to an electronic interface board (Neuralynx Inc., Bozeman, MT) and bent at 4.5mm from the tip to maximize the proximity of the fUSI probe to the skull (Fig. 1B). Recordings took place over one hour, animals were injected after 5 minutes with saline vehicle or MK-801 (1mg/kg) and stimulation or sham stimulation was delivered for 5 minutes 45 minutes into the recording period (Fig. 1C). All procedures were approved by the University of Southern California, Institutional Animal Care and Use Committee (IACUC #21006).

### Histology

Mice were euthanized immediately after the fUSI recording by isoflurane overdose followed by transcardial perfusion using 50 mL of 0.1M sodium phosphate buffer saline (PBS) and 50 mL of 4% paraformaldehyde. Brains were harvested and stored in phosphate buffered saline at 4^°^C. To confirm electrode positioning within the MSN, coronal sections were cut at 100µm thickness with a vibratome (Leica VT 1200; Leica Biosystems, Buffalo Grove, IL) and then Nissl stained with Cresyl Violet (Fig. 2A,B).

**Figure 2.**
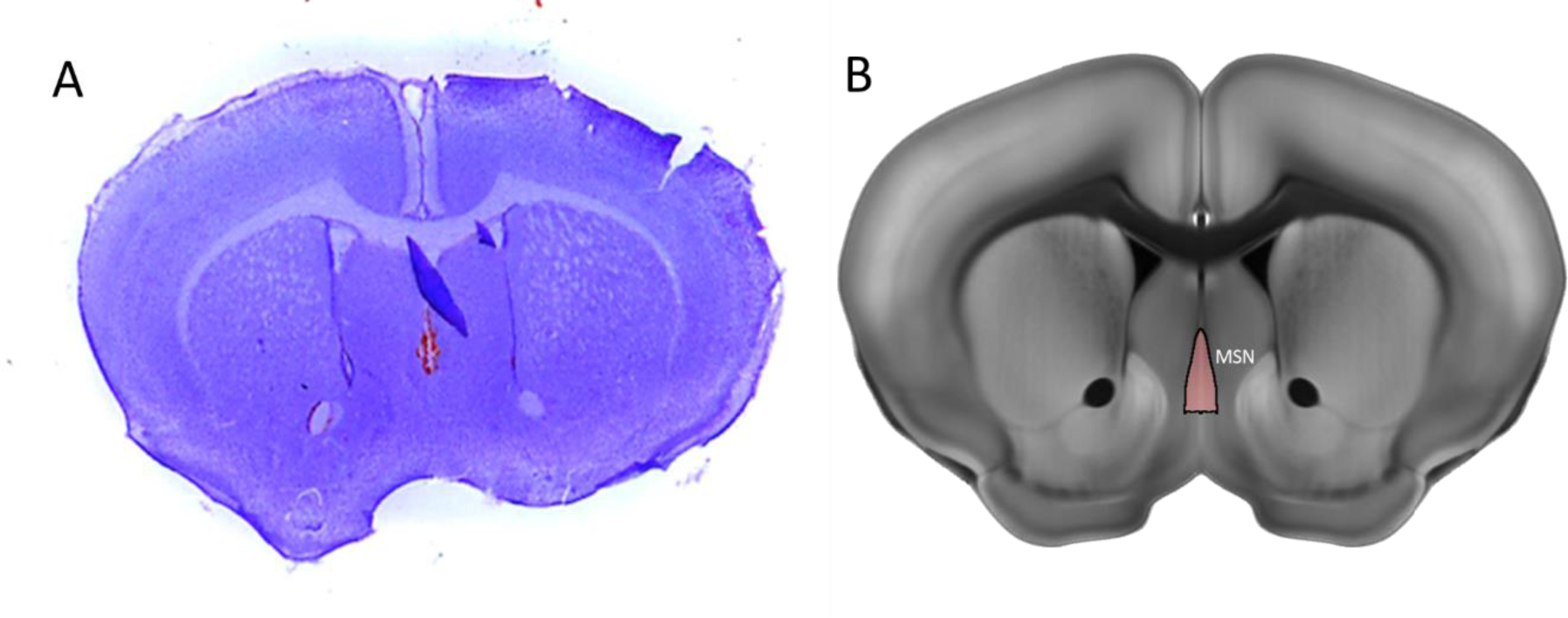
Histological mapping of electrode placement in the MSN. **A)** Representative Nissl stain of the mouse brain with a blood mark indicating the electrode placement **B)** Annotation of the MSN from the Allen Reference Atlas – Mouse Brain in the same slice position as A.

### Computational Modeling of the Induced Voltages: Volume of Tissue Activation

Our group has developed a computational analysis method for applications in electrophysiology and bioelectromagnetic interactions, namely the Admittance Method (AM)[32]. AM discretizes a bulk tissue model into cuboid voxels, where each voxel is represented by an equivalent circuit of lumped passive elements. The admittance value at each voxel is calculated using the material properties such as conductivity and permittivity. A set of linear equations using iterative methods are used to calculate the induced voltage at each node of the voxelized network. Here, a 3-D bulk tissue model with the desired electrode configuration was constructed, a current-controlled stimulation pulse was applied through the electrodes and the induced voltages at each voxel were calculated. A mouse head model (Fig. 3A) was used for this study where three types of tissue were considered (grey matter, white matter (WM) and cerebrospinal fluid (CSF). Electrodes were set as platinum and the medium surrounding the mouse head was set as air. The locations and geometries of the WM and CSF with respect to the septal medial nucleus were approximated using an interactive online mouse brain atlas (http://labs.gaidi.ca/mouse-brain-atlas/). The output from AM was then processed using a MATLAB script to produce an illustration of the voltage mapping in the mouse head (Fig. 3B). A one-dimensional voltage profile in the proximity of the electrodes was plotted to estimate the volume of tissue that was electrically affected/activated (Fig. 3C).

**Figure 3.**
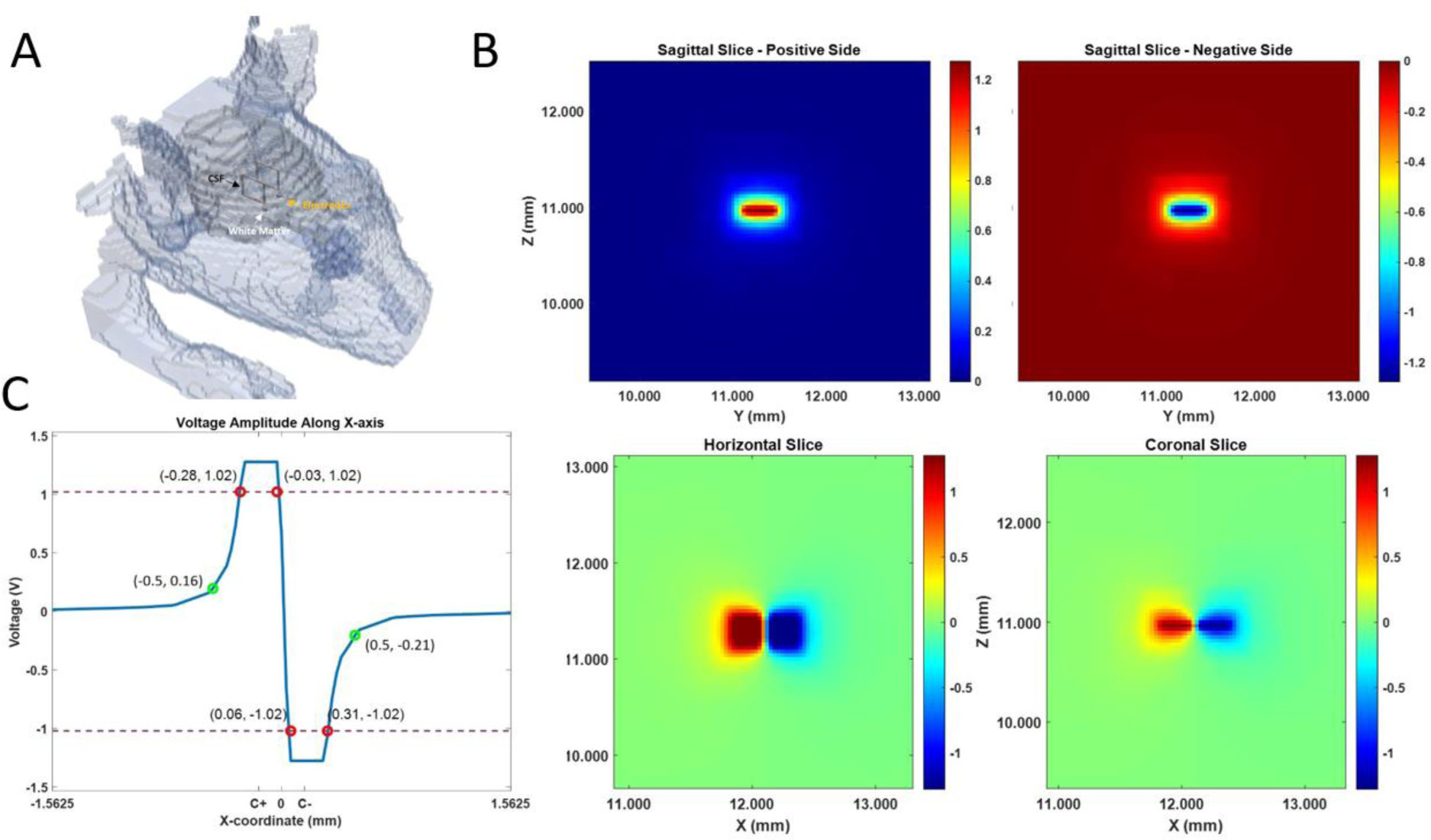
Volume of Tissue Activation. **A)** 3-D illustration of the discretized mouse head model. The bulk tissue model is divided into voxels at a resolution of 31.25 μm. Each voxel represents an equivalent electrical circuit according to its material type. The brain model is mostly uniform as grey matter. The proximity of the medial septal nucleus, where the electrodes are placed, also contains areas of white matter and CSF modeled as basic geometries. **B)** The voltage mapping near the medial septal nucleus when stimulated with a bipolar electrode configuration. Top row: two sides of the sagittal slice, looking into both the positively and negatively polarized electrodes. Bottom row: the horizontal and coronal slices capturing the voltage distribution around both electrodes. The absolute voltage decays to zero rapidly moving away from the electrodes, demonstrating that the volume of activation is confined within a diameter of about 1 millimeter. **C)** The voltage amplitude plotted along a single axis. The voltage decay profile moving away from the electrode is very similar along all three axes and the volume of activation can be modeled as a sphere. The “equivalent diameter” for the volume affected by stimulation is approximated by choosing points where the slope is approaching zero (green circles), which determines the current density generated at that location. Thresholding can also be used to visualize the points where the target voltage amplitude is maintained. The red circles label the coordinates at which an amplitude of 1 Volt or more is observed. C+ and C-labels mark the centers of the positive and negative electrodes, respectively.

### Data analysis

#### Data pre-processing

The Iconeus One acquisition system generates power Doppler (pD) images pre-processed with built-in phase-correlation based sub-pixel motion registration and singular-value-decomposition (SVD) based clutter filtering algorithms [33]. These algorithms were used to separate tissue signal from blood signal to obtain pD images (Fig. 4A/B). To correct potential physiological and motion artifacts, we adopted rigid motion correction techniques that have successfully been used in fUSI and other neuroimaging studies [34–36]. These were combined with high frequency filtering algorithms to eliminate noise artifacts. We employed a low-pass filter with normalized passband frequency of 0.02 Hz, with a stopband attenuation of 60 dB that compensates for delay introduced by the filter, to remove high-frequency fluctuations in the pD signals.

**Figure 4.**
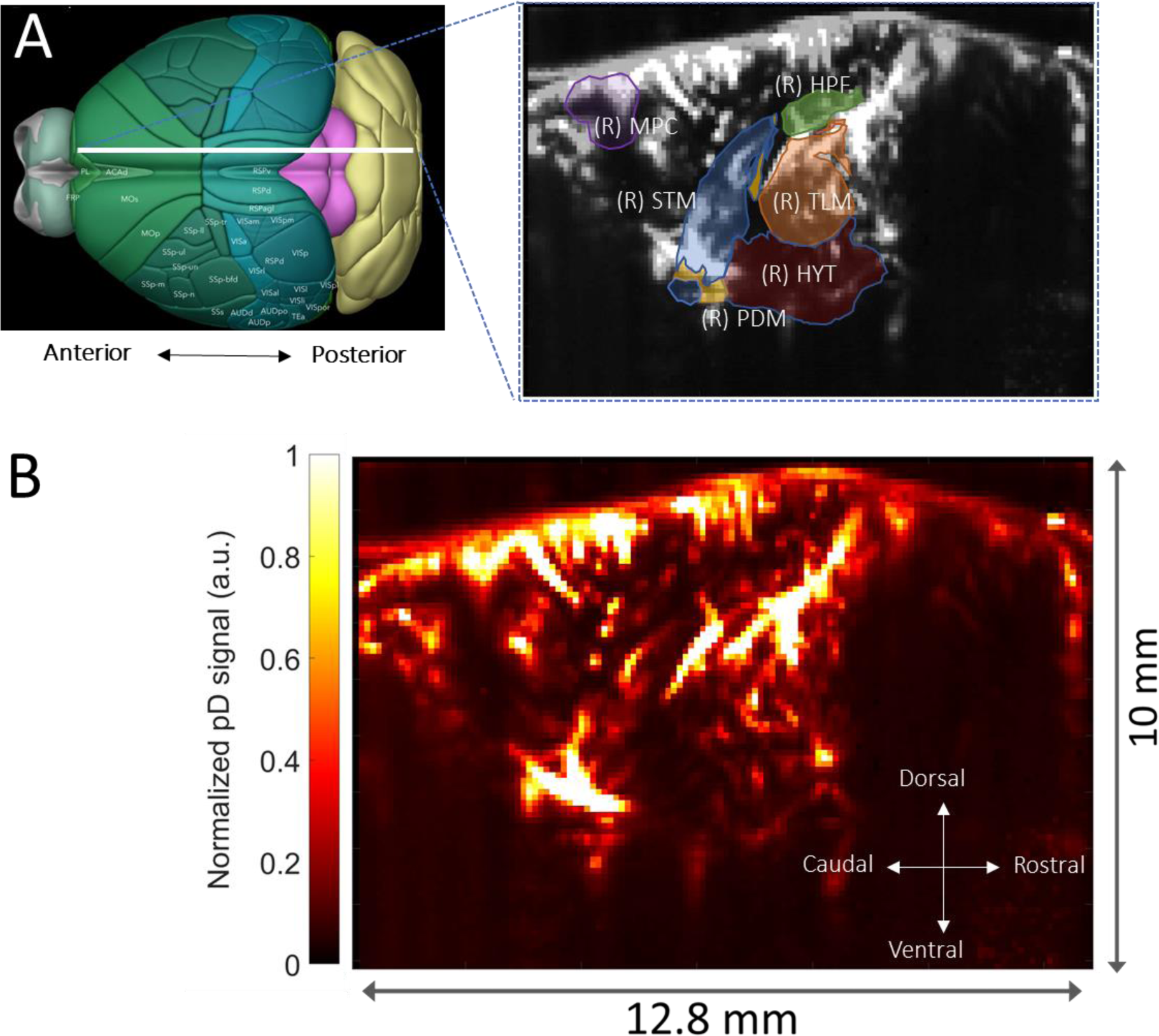
Functional ultrasound imaging (fUSI) of the mouse brain. **A)** 3D mouse brain model with fUSI probe positioning (white bar) and ROIs – hippocampus (HPF), medial prefrontal cortex (mPFC), hypothalamus (HYT), thalamus (TLM), pallidum (PDM), and striatum (STM), superimposed onto a mean grayscale fUSI vascular map of the sagittal mouse brain. **B)** pD image of cerebral blood volume (CBV) in a sagittal plane (max-min normalized relative scale).

#### Effects of MK-801 on CBV

We investigated the temporal effects of intraperitoneal MK-801 administration on the septohippocampal circuit (hippocampus, mPFC) and surrounding regions (striatum, pallidum, thalamus, hypothalamus). To do so, we generated event-related average (ERA) time course curves of the CBV changes (ΔCBV) as a percentage change of the pD signal from baseline activity for the selected ROIs. The average pD signal from 2 minutes prior to the saline or drug injection was used as the baseline. We utilized a repeated measures analysis of variance (rmANOVA) to assess the effects and interactions between drug (saline and MK801) and ROI factors over time. We fitted a repeated measures ‘*within-design’* model to the CBV percentage change signals over 42-min interval (including 2 minutes just before the drug injection and the 40 minutes after injection) for each mouse and ROI for the rmANOVA analysis. To further quantify the relative differences in ΔCBV between saline-vehicle and MK-801-treated mice in various ROIs, we used the last 2 minutes of recordings to compute the mean effects-size differences in ΔCBVs and the 95% confidence interval of the effect size (if 95% confidence interval contains zero, then the effect is not significant at the 0.05 level) in each ROI. We also computed the Cohen’s *d* value in each ROI as a measure of the drug effect size that describes the standardized difference between the means of ΔCBVs in the two groups of animals [37]. A Cohen’s d value of 0.2 represents a small effect size, 0.5 represents a moderate effect size, 0.8 represents a large size and greater than 0.8 represents a very large size.

#### Effects of MSN DBS during stimulation on brain hemodynamics

We computed the ΔCBV for each ROI to investigate effects of MSN theta- and gamma-frequency stimulation on the cerebral hemodynamics following saline control or MK-801 injection. We observed ERA time-series of the ΔCBV of each selected ROI relative to the average pD signal acquired 2 minutes prior to stimulation onset to visualize the temporal dynamics of DBS effects to CBVs of the ROIs (Fig. 1C). We utilized a three-way rmANOVA to assess the effects and interactions between drug (saline and MK801), stimulation (gamma, theta, no-stimulation) and ROI factors across time during the stimulation process. Here, we fit a repeated measures ‘*within-design’* model to the ΔCBV signals over the 7-min period (including 2 minutes prior to stimulation, 5 minutes during stimulation) for each animal and ROI. Subsequently, we computed the mean effects-size differences in ΔCBVs, the 95% confidence interval of the effect size, and the Cohen’s *d* value in each ROI as a measure of the stimulation effect size. We measured the mean ΔCBVs during stimulation in theta [saline: n=16, MK-801: n=12], gamma [saline: n=14, MK-801: n=14], and no-stimulation [saline: n=14; MK-801: n=10] animal groups. The mean ΔCBVs were calculated utilizing the last 2 minutes of pD signal during stimulation across animals in each of the three stimulation categories.

#### Effects of MSN DBS after stimulation offset on brain hemodynamics

We are also interested in assessing the effects of MSN DBS after stimulation offset. To do so, we repeated the same analysis described above, but we used the 10 minutes of pD signal after stimulation offset, plus 2 minutes prior to stimulation onset as a baseline. The mean ΔCBVs were calculated utilizing the last 2 minutes of pD signal after stimulation offset across animals in each of the three stimulation categories. Our goal is to identify whether MSN DBS causes persistent and delayed changes in CBVs within and/or outside the septohippocampal network.

#### Statistical analysis of drug and stimulation effects on ΔCBVs

All the analysis was performed using Matlab Version 9.13.0.2193358 (R2022b). We assessed the drug and stimulation effects and interactions utilizing the Matlab functions *‘fitrm’* and *‘ranova’* for the repeated measures model fitting and the rmANOVA respectively. We utilized the Greenhouse-Geisser approximation to correct for the possibility of non-compound symmetry (same variance in means and shared common correlation in paired responses) in the ROI-time series assessed. The mean effect-size differences and the *Cohen’s d* values for the mean difference in ΔCBVs between saline and MK801, as well as the between no-stimulation-gamma, no-stimulation-theta, and gamma-theta were computed with the *‘meanEffectSize’* Matlab function.

## Results

### NMDA antagonist MK-801 reduced blood perfusion

We analyzed 40 minutes of pD signal from the septohippocampal circuit (hippocampus, mPFC) as well as surrounding structures (hypothalamus, thalamus, pallidum, and striatum) to assess the effects of MK-801 on cerebral hemodynamics (Fig. 4A). We quantified changes in CBV (ΔCBV) as a percent change relative to baseline activity (average of 2 minutes pD signal acquired prior to drug injection). A repeated measures ANOVA (factors treatment × ROI; where treatment is saline vs. MK-801 and ROI is the 6 recorded brain areas) revealed that there was a statistically significant effect over time (F (2519, 56406) = 8.76, p = 6.3 e-5) after Greenhouse-Geiser approximation correction. To quantify the effect of MK-801 on CBV, we computed the percentage change (i.e., ΔCBV) relative to baseline using the last two minutes before stimulation onset (38-40 minutes after drug injection) and found a reduction of ΔCBV (mean ± SEM) in the hippocampus (-3.6 ± 1.1 %), mPFC (-4.1 ± 0.61 %), hypothalamus (-1.0 ± 0.2 %), pallidum (-1.5 ± 0.4 %), striatum (-1.7 ± 0.3 %), and thalamus (-3.0 ± 0.5 %). For saline-treated animals these values were: hippocampus (-0.7 ± 0.6 %), mPFC (-0.1 ± 0.2 %), hypothalamus (-0.7 ± 0.6 %), pallidum (0.9 ± 0.4 %), striatum (-0.3 ± 0.4 %), and thalamus (0.2 ± 0.4 %) (Fig. 5A-F, radar chart insert). The mean effect-size difference in ΔCBVs and Cohen’s d analysis comparing mean ΔCBV induced by saline and MK-801 over a 2-minute interval (38 – 40 minutes post-drug injection), revealed that MK-801 induces greater decreases in CBV than saline control in all ROIs investigated (Fig. 5A-F, radar chart insert) - i.e., mPFC (ΔCBV mean difference between saline and MK-801 animals ± confidence, Cohen’s d; 3.96 ± 0.38 %, d= 0.42), thalamus (3.17 ± 0.23 %, d= 0.55), hippocampus (2.82 ± 0.42 %, d= 0.27), pallidum (2.42 ± 0.20 %, d= 0.49), hypothalamus (1.72 ± 0.12 %, d= 0.59) and striatum (1.33 ± 0.18 %, d= 0.29). Together, these results indicate that MK-801 reduces CBV both within and outside of the septohippocampal network.

**Figure 5.**
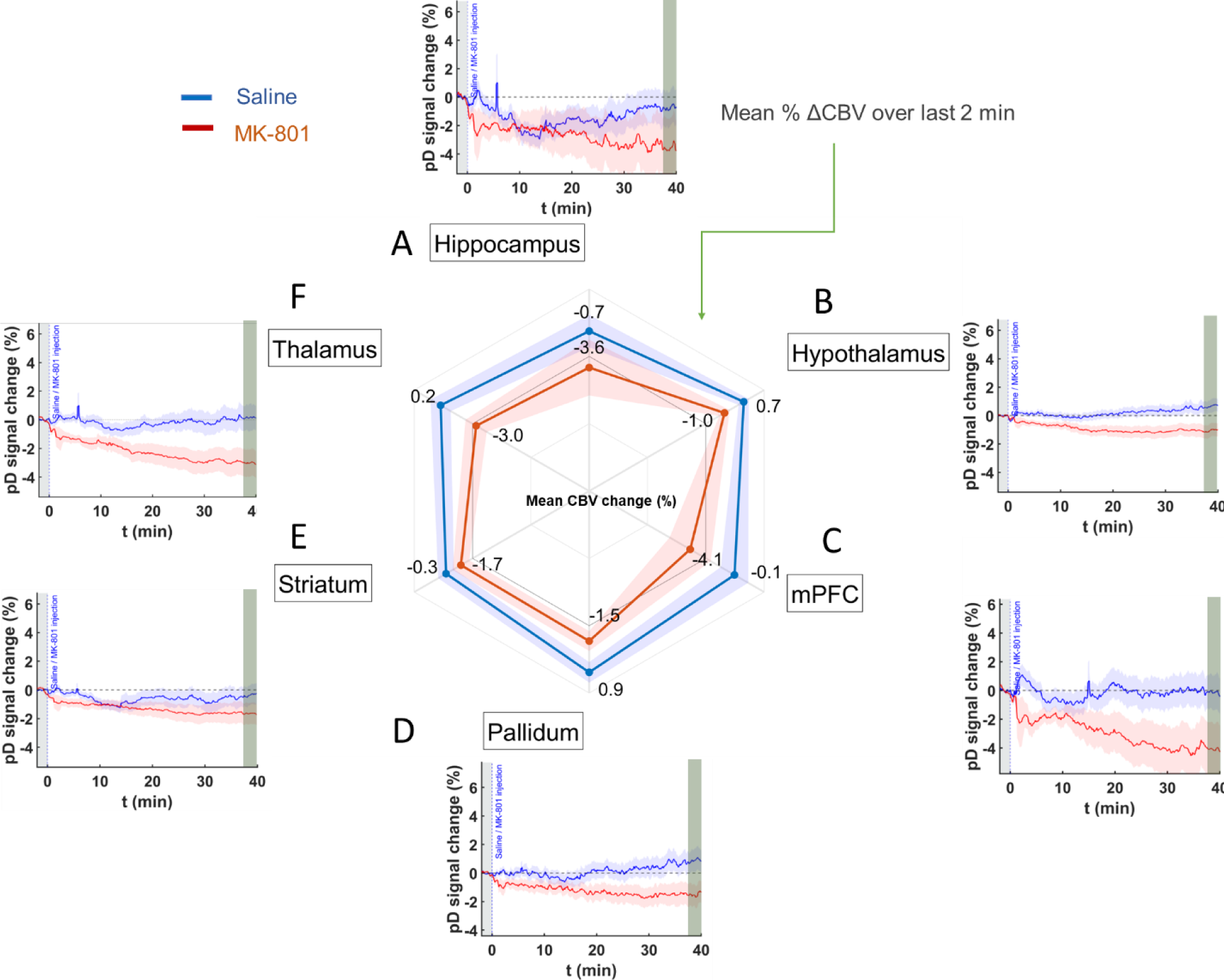
Event related average (ERA) temporal course curves prior to stimulation onset and mean ΔCBVs during the last 2 minutes in saline and MK-801 treated animals. **A – F)** Temporal course (42 minutes) of ΔCBV relative to baseline (2-minute average pD signal before saline or MK-801 drug injection) in the **A)** hippocampus, **B)** hypothalamus, **C)** mPFC, **D)** pallidum, **E)** striatum, and **F)** thalamus after saline [blue] and 1.0 mg/kg MK-801 [1] injection. The radar chart insert shows MK-801-induced decreases in CBV in all ROIs compared to saline over the last 2 minutes interval from 38-40 minutes post injection.

### MSN stimulation increases CBV in saline control animals in a frequency- and region-dependent manner

We assessed whether theta- and gamma-frequency MSN stimulation has disparate impacts on CBV measures in saline-treated control mice. ERAs of ΔCBV reflect the temporal responses to theta, gamma, and no stimulation during and post stimulation time periods (Fig. 6, 7). A three-way repeated measures ANOVA (factors, treatment × ROI × DBS; where treatment is saline vs. MK-801, ROI is the 6 recorded brain areas, and DBS is theta frequency vs. gamma frequency vs no DBS) was utilized to examine the effects and interactions of drug, stimulation, and ROIs during the 5 minutes period after onset of MSN stimulation. The two-minute baseline period pre-stimulation was included in the analysis. We found significant effects of drug over time (F(419, 186036) = 7.35, p = 1.85 e-8), stimulation over time (F(823, 186036) = 2.71, p = 7.20 e-4), as well as, interaction of drug and stimulation over time (F(838, 186036) = 1.96, p = 1.95 e-2) during the 5 minutes stimulation interval, after Greenhouse-Geisser approximation correction. To further quantify the effects of DBS, we computed the mean ΔCBV in the last 2 minutes during stimulation across animals in each stimulation category and compared the mean-effect size differences in ΔCBVs between no stimulation and stimulation in each ROI. We found that MSN theta stimulation increased CBV compared to no-stimulation only in the mPFC (mean ΔCBV difference between theta- and no-stimulation ± confidence, Cohen’s d; 0.82 ± 0.12 %, d= 0.45) and hippocampus (0.39 ± 0.12 %, d= 0.21). For the rest on ROIs the effect size magnitude was either very small (i.e., Cohen’s d <0.08) or theta-frequency stimulation caused further reduction in CBVs compared to no-stimulation. On the other hand, MSN gamma stimulation caused increases in CBV compared to no-stimulation in the mPFC (0.60 ± 0.11 %, d= 0.36), pallidum (0.38 ± 0.09 %, d= 0.30) and striatum (0.22 ± 0.06 %, d= 0.26). For the rest of the ROIs the effect size magnitude was either very small (i.e., Cohen’s d < 0.095) or gamma-frequency stimulation resulted in further reduction in CBVs compared to no-stimulation. When comparing the ΔCBV induced by the theta and gamma stimulation in mPFC – the only ROI that exhibited moderate effect on both types of stimulations – we found very small effect size on ΔCBV between the two types of stimulations (mean ΔCBV differences between theta- and gamma-stimulation ± confidence, Cohen’s d; 0.22 ± 0.10, d= 0.14).

**Figure 6.**
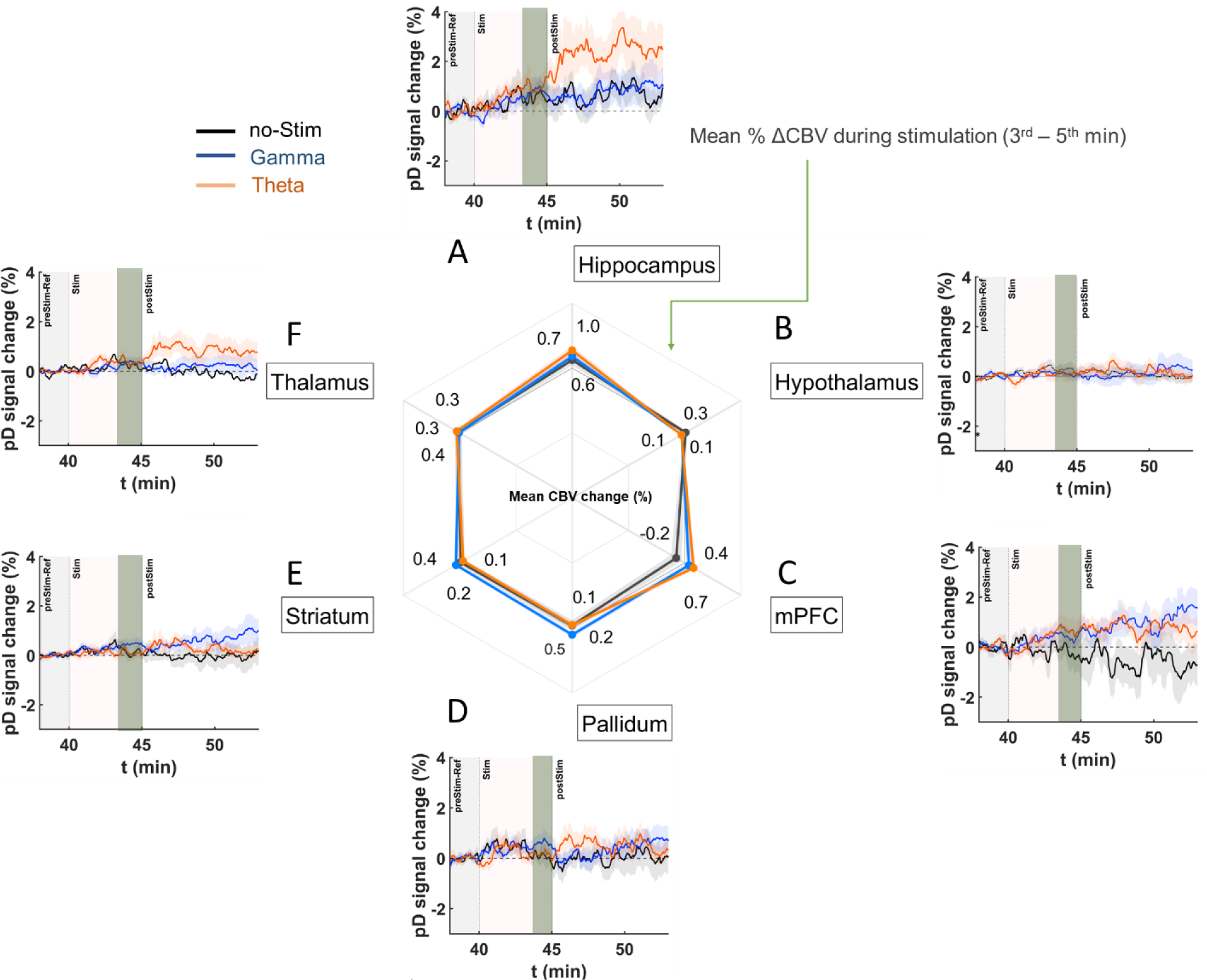
DBS ERA curves after stimulation onset and mean ΔCBVs in saline mice during the last 2 minutes of stimulation period. **A – F)** Temporal course (theta [orange], gamma [blue], no-stimulation [black]) of mean ΔCBV relative to baseline (2 minutes average pD signal prior to DBS) for the **A**) hippocampus, **B**) hypothalamus, **C**) mPFC, **D**) pallidum, **E**) striatum, and **F**) thalamus regions in the saline-treated animals. Radar chart insert gives the mean percentage ΔCBVs during stimulation for theta [orange], gamma [blue], and no-stimulation [dark gray] animals in the ROIs investigated. Means were calculated utilizing the last 2 minutes of pD signals acquired during stimulation (3^rd^ – 5^th^ minute after stimulation onset) across animals in each stimulation category.

The next step was to assess the effects of DBS after stimulation offset (i.e., post-stimulation). To do so, we conducted a three-way repeated measures ANOVA (factors, treatment × ROI × DBS; where treatment is saline vs. MK-801, ROI is the 6 recorded brain areas, and DBS is theta frequency vs. gamma frequency vs no DBS) over the 10 minutes period after the offset of MSN stimulation. We found significant effects of drug over time (F(1019, 452436) = 5.28, p = 1.27 e-4), stimulation over time (F(2038, 452436) = 3.67, p = 1.17 e-4), as well as, interaction of drug and stimulation over time (F(2038, 186036) = 3.09, p = 9.40 e-4), after Greenhouse-Geisser approximation correction, across the 10 minutes post-stimulation interval. We further quantified the post-effects of DBS by computing the mean ΔCBV in the last 2 minutes of the acquisition – i.e., 8-10 minutes post-stimulation and comparing the mean-effect size differences on ΔCBV between no stimulation and stimulation in each ROI. The results showed that MSN theta stimulation causes increases in CBV compared to no-stimulation in the hippocampus (mean ΔCBV differences between theta- and no-stimulation ± confidence, Cohen’s d; 1.30 ± 0.21, d= 0.42), mPFC (1.20 ± 0.22 %, d= 0.37) and thalamus (0.97 ± 0.11 %, d= 0.58) (Fig. 7A, F – radar chart). On the other hand, MSN gamma stimulation resulted in an increase in CBV compared to no stimulation in the mPFC (2.01 ± 0.22 %, d= 0.60), striatum (0.77 ± 0.14 %, d= 0.37) and pallidum (0.61 ± 0.16 %, d= 0.25). When comparing the differences in ΔCBVs induced by theta and gamma stimulation in the mPFC – the only ROI that exhibited medium to moderate effect on both types of stimulations – we found that gamma induces higher ΔCBV than theta stimulation with medium effect size difference (mean ΔCBV differences between gamma- and theta-stimulation ± confidence, Cohen’s d; 0.81 ± 0.17, d= 0.31).

**Figure 7.**
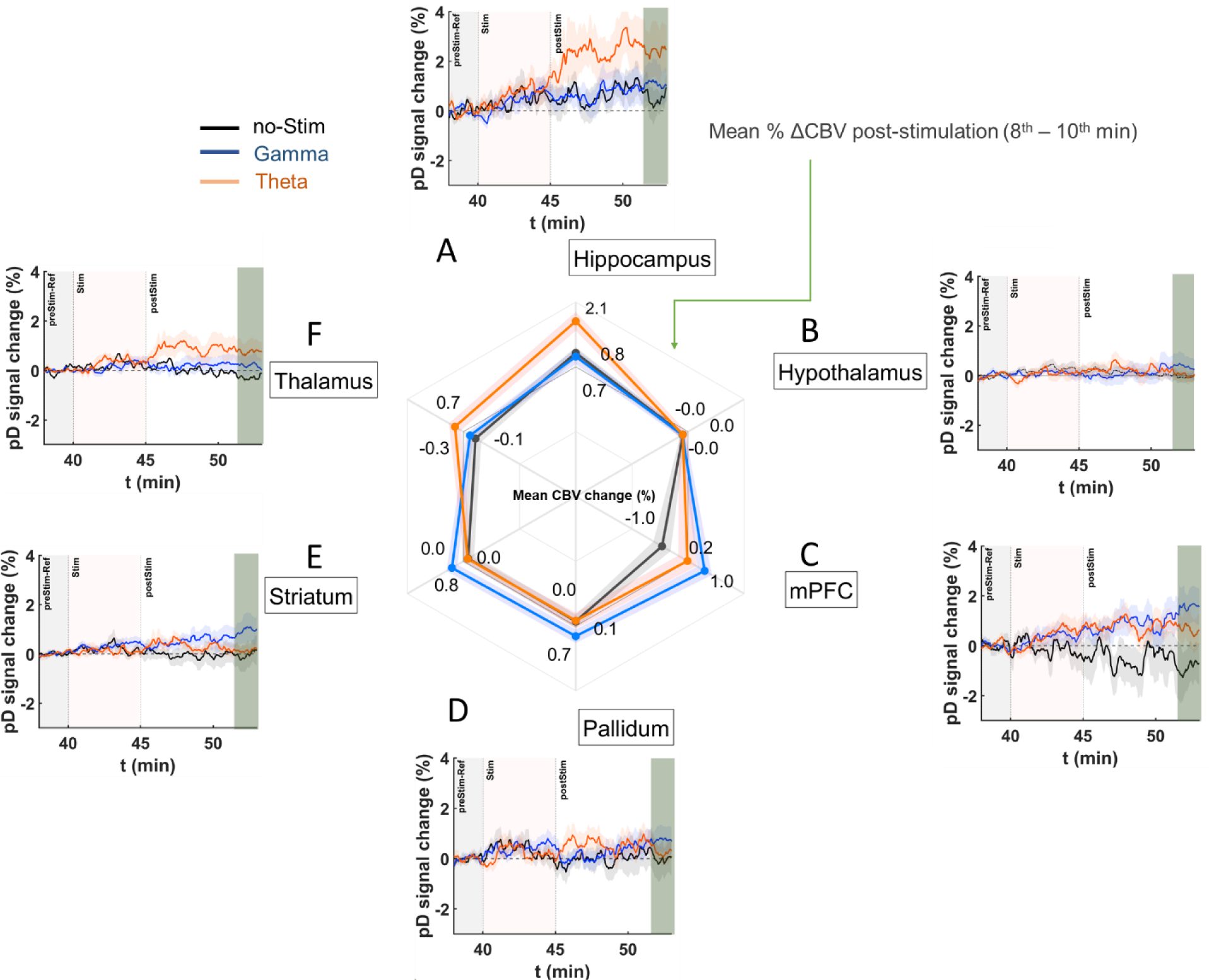
DBS ERA curves after stimulation onset and mean ΔCBVs in saline mice during the last 2 minutes of recordings in the post-stimulation period in 6 ROIs. Similar to Figure 6, but the radar chart gives the mean percentage ΔCBVs during post-stimulation for theta [orange], gamma [blue], and no-stimulation [dark gray] for the **A)** hippocampus, **B)** hypothalamus, **C)** mPFC, **D)** pallidum, **E)** striatum, and **F)** thalamus regions in the saline-treated animals. Means were calculated utilizing the last 2 minutes of pD signals acquired post stimulation (8^th^ – 10^th^ minute after stimulation offset) across animals in each stimulation category.

### Theta-frequency stimulation elicits stronger CBV increases than gamma-frequency stimulation in MK-801 treated animals

Recently, our group showed that theta, but not gamma frequency DBS of the MSN improves spatial memory in MK-801 treated rats [27]. Therefore, we sought to determine if MSN theta and gamma frequency stimulation had differing impacts on neurovascular activity measures within memory-associated regions including the mPFC and hippocampus as well as neighboring regions outside the septohippocampal network (striatum, pallidum, thalamus, hypothalamus) following MK-801 drug-administration. Again, we assessed the mean effect-size differences in ΔCBV between MSN theta-, gamma-, and no-stimulation in each of the selected ROIs, relative to 2 minutes of pD signal recordings just prior to stimulation onset. The analysis was performed after repeated measures ANOVA over the stimulation and post-stimulation time intervals to examine the effects and interactions of drug, stimulation, and ROI (results presented in previous section). Fig. 8 and 9 display the ERA curves for ROIs in response to theta, gamma, and no DBS during and post stimulation periods in the MK-801 group. We found that MSN theta stimulation in the MK-801 treated group caused increased ΔCBV with respect to no-stimulation group with medium to large effect size in all ROIs except mPFC – i.e., hippocampus (mean ΔCBV differences between theta- and no-stimulation ± confidence, Cohen’s d; 2.01 ± 0.20 %, d= 0.78), thalamus (0.49 ± 0.07 %, d= 0.54), pallidum (0.36 ± 0.07 %, d= 0.43), striatum (0.18 ± 0.04 %, d= 0.31) and hypothalamus (0.14 ± 0.04 %, d= 0.30) (Fig. 8). Intriguingly, we only found a medium effect-size increase in ΔCBV for the pallidum (0.60 ± 0.08 %, d= 0.53) and striatum (0.23 ± 0.05 %, d= 0.38) between MSN gamma-stimulation and no-stimulation groups, during the stimulation period. For the rest of the ROIs, the effect-size was very small.

**Figure 8.**
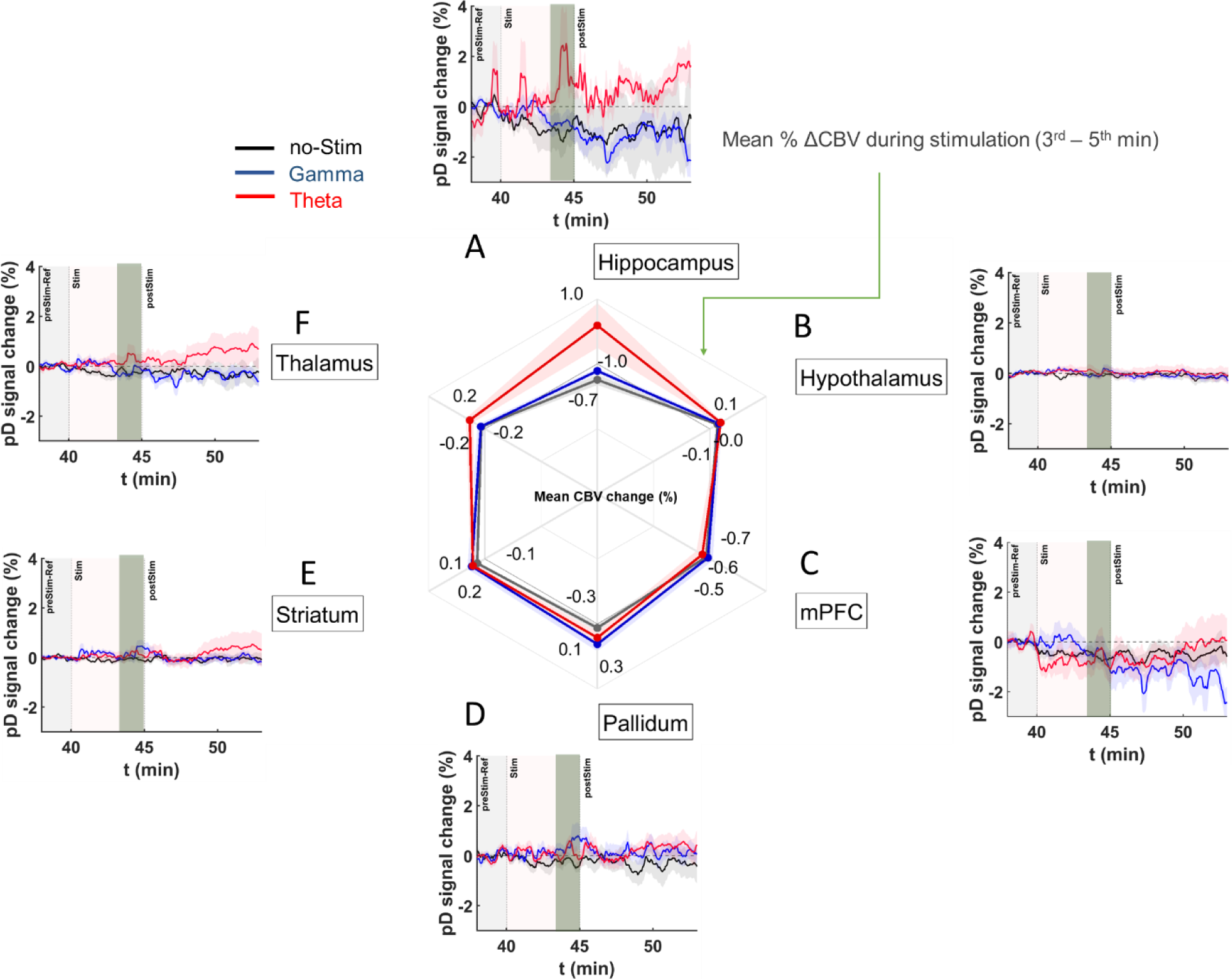
DBS ERA curves after stimulation onset and mean ΔCBVs in MK-801 treated mice during the last 2 minutes of stimulation period. **A – F)** Temporal course (theta [1], gamma [blue], no-stimulation [black]) of mean ΔCBV relative to baseline (2 minutes average pD signal prior to DBS) for **A)** hippocampus, **B)** hypothalamus, **C)** mPFC, **D)** pallidum, **E)** striatum, and **F)** thalamus regions in the MK-801 drug injected mice. Radar chart insert gives the mean percentage ΔCBVs during stimulation for theta [1], gamma [blue], and no-stimulation [dark gray] animals in the ROIs investigated. Means were calculated utilizing the last 2 minutes of pD signals acquired during stimulation (3^rd^ – 5^th^ minute after stimulation onset) across animals in each stimulation category.

**Figure 9.**
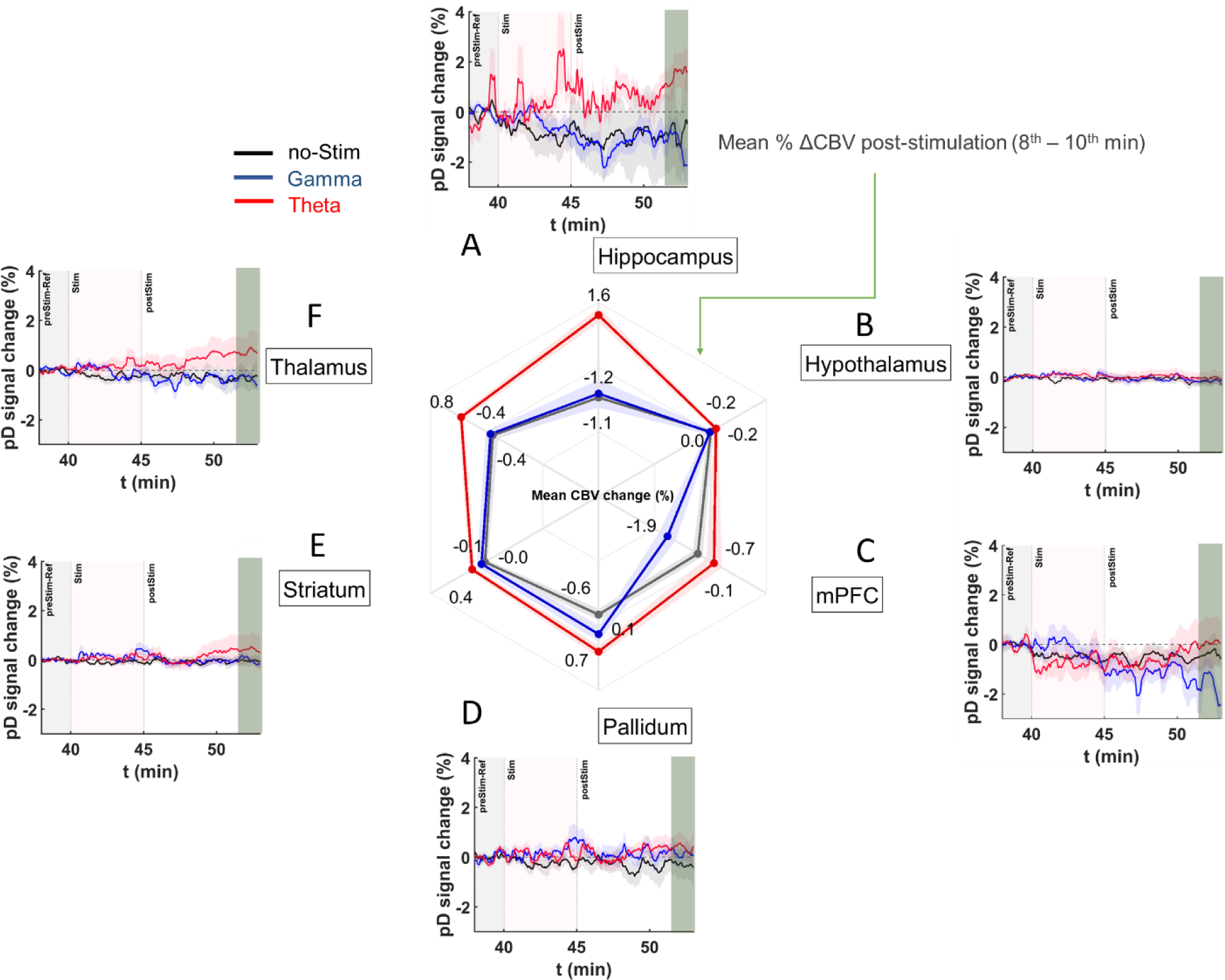
DBS ERA curves after stimulation onset and mean ΔCBVs in MK-801 treated mice during the last 2 minutes of recordings in the post-stimulation period. Similar to Figure 8, but the radar chart gives the mean percentage ΔCBVs in post-stimulation period for theta [1], gamma [blue], and no-stimulation [dark gray] for the **A)** hippocampus, **B)** hypothalamus, **C)** mPFC, **D)** pallidum, **E)** striatum, and **F)** thalamus regions in the MK-801 treated animals. Means were calculated utilizing the last 2 minutes of pD signals acquired post stimulation (8^th^ – 10^th^ minute after stimulation offset) across animals in each stimulation category.

Importantly, we found that ΔCBV increased in theta-relative to no-stimulation animals after stimulation offset with medium to larger effects in the pallidum (1.26 ± 0.15 %, d=0.66), hippocampus (2.8 ± 0.37 %, d=0.60) and thalamus (1.24 ± 0.17 %, d= 0.57) and small to medium effects in the striatum (0.50 ± 0.13 %, d= 0.31), hypothalamus (0.21 ± 0.06 %, d= 0.28) and mPFC (0.64 ± 0.22 %, d= 0.22) (Fig. 9). Importantly, we found only small to medium effects of gamma MSN stimulation on ΔCBV in the pallidum, relative to stimulation animals, and after stimulation offset (mean ΔCBV differences between gamna- and no-stimulation ± confidence, Cohen’s d; 0.66 ± 0.13 %, d=0.39) (Fig. 9). For the rest of the ROIs, the effect-size was either very small (i.e., Cohen’s d < 0.1 in hippocampus, striatum and thalamus) or gamma-frequency stimulation resulted in further CBV reduction (i.e., in hypothalamus and mPFC) compared to no-stimulation.

Additionally, our results showed medium or large mean effect-size differences in ΔCBVs between the theta and gamma stimulated MK801 groups after stimulation offset in the hippocampus (2.67 ± 0.28 %, d= 0.70), mPFC (1.83 ± 0.23 %, d= 0.59), and thalamus (1.15 ± 0.14 %, d= 0.56) (Fig. 9). Together these results demonstrate that theta-frequency stimulation elicits the strongest CBV response in MK-801-treated mice in the hippocampus and pallidum, while gamma stimulation had almost no effect on hippocampal CBV (Cohen’s d = 0.03) and decreased CBV in the mPFC compared to no-stimulation mice.

## Discussion

The present study utilized the high spatiotemporal resolution and sensitivity of fUSI to demonstrate that acute administration of MK-801 causes a significant reduction in CBV across all ROIs. Furthermore, we demonstrated that theta frequency MSN DBS alters regions within the septohippocampal network, with the strongest effect on the hippocampus. Intriguingly, the observed increase in hippocampal CBV remain even after cessation of DBS. On the other hand, structures outside the septohippocampal network, such as the hypothalamus and striatum, show less of a response to DBS. These effects were less pronounced with gamma frequency stimulation with very small effects on the hippocampus. These findings suggest that MSN theta frequency DBS precisely modulates neurovascular activity in cognitive networks [27].

### MK-801 reduced CBV in all ROIs

MK-801 and other NMDA antagonists have been widely used in preclinical models to mimic the behavioral and electrophysiological deficits associated with schizophrenia [16,18,38,39]. However, the regionally specific effects of MK-801 on CBV in such models is not well known. We observed that MK-801 reduced CBV across all ROIs. Importantly, previous fMRI studies have observed reduced BOLD signals in hippocampal and prefrontal areas in schizophrenia patients [40–42]. In this context, our findings support the use of MK-801 as a neurovascular model of schizophrenia. Furthermore, our study demonstrates the feasibility of using fUSI to identify network-specific hemodynamic changes as an additional modality for studying neurocognitive disorders.

### MSN theta stimulation was relatively specific to cognitive networks

We observed that theta frequency MSN DBS resulted in an increase to CBV in the hippocampus in both the saline- and MK-801-treated animals. Importantly, this effect was greatest in the hippocampus, which receives direct projections from the MSN, and is a primary target for neuromodulatory interventions to treat memory dysfunction [23–25,27]. In saline control animals, significant increases to CBV were also observed in the mPFC (during and after stimulation) and thalamus (after stimulation), both of which play important roles in memory function and receive direct projections from the MSN (Fig. 2A) [22]. Interestingly, gamma stimulation did not alter hippocampal CBV, but did increase regions of the brain that were anatomically closer to the MSN. Specifically, mPFC CBV was increased in saline-treated animals, while the pallidum and striatum were increased to a lesser degree in MK-801-treated animals. This suggests that MSN gamma stimulation may have a local response to stimulation but is less specific to the neural circuitry being stimulated.

### MSN theta stimulation increased hippocampal CBV during and after stimulation despite NMDA antagonism

A leading hypothesis is that reduced N-methyl-D-aspartate (NMDA) receptor-mediated glutamatergic transmission underlies psychiatric conditions involving cognitive and memory dysfunction [16,18,38,39,43,44]. Reductions to NMDA activity either pharmacologically or through genetic manipulation have been shown to decrease theta activity, increase gamma activity and lead to deficits in spatial navigation and memory [16,17,27,45,46]. Research by our group has found that acute (<5 minutes) theta frequency (7.7 Hz), but not gamma frequency (100 Hz) stimulation of the MSN during the Barnes maze task improves spatial memory in rodents following pharmacological NMDA antagonism [27]. Further, we have also demonstrated that MSN theta stimulation prior to the task can also improve spatial memory [23]. Interestingly, hippocampal theta oscillations return to baseline after cessation of MSN DBS. Therefore, the question remains open as to how MSN DBS mediates sustained improvements. In this current study, we demonstrate that hippocampal CBV remains elevated after cessation of MSN stimulation.

### MSN theta stimulation may drive high frequency or spiking activity via hippocampal interneurons

A number of studies utilizing various modalities in combination with fUSI have found a strong relationship between pD signal and neuronal activity [47–49]. This relationship was also true for high frequency oscillatory activity (∼100Hz) but was much weaker for lower frequency oscillations [49]. This suggests that the theta-induced increases to hippocampal CBV may reflect increased hippocampal gamma or spiking activity, rather than increases to theta oscillatory activity itself. However, given that gamma band activity is often correlated with spiking activity, differentiating between changes in oscillatory dynamics and spiking activity is not possible in the present study [50].

GABAergic interneurons in the hippocampus play an important role in synchronizing hippocampal oscillatory activity. Indeed, inhibitory neurons are hypothesized to be a primary source of dysfunction in pathologies involving NMDA dysfunction (for review see [51]) and are inhibited by NMDA-antagonists [52,53]. One possibility is that theta frequency MSN stimulation may, by briefly stimulating afferent populations at theta-frequency, act as a ‘reset’ and allow synchronous innervation of hippocampal interneurons that regulate the activity of glutamatergic pyramidal cells. Indeed, it has been previously shown that optogenetic stimulation of GABAergic neurons decreases spontaneous neural activity and leads to an increase in local blood flow [54]. This hypothesis is further supported by a recent study by Nunez-Elizadle and colleagues who observed that the relationship between the fUSI signal and firing rates were greatest for putative interneurons [49]. Future studies combining single unit activity and fUSI could test this hypothesis.

### MSN gamma stimulation did not affect hippocampal CBV

Our previous work suggests that MSN theta, but not gamma stimulation can improve spatial memory in MK-801-treated rodents [27]. While MSN theta stimulation increased hippocampal CBV during and after stimulation in MK-801-treated animals, this was not true of MSN gamma stimulation. Gamma stimulation resulted in delayed increases to mPFC CBV in saline-treated animals and had no effect on MK-801-treated animals in any of the ROIs. These results suggest that MSN gamma stimulation is not sufficient to engage hippocampal activity and highlights the importance of frequency parameters in DBS paradigms for spatial memory. This is further supported by our previous study demonstrating no improvement to spatial memory in MK-801 treated animals with MSN gamma stimulation [27].

### Implications for neuromodulation

We observed that theta-frequency stimulation of the MSN increased blood perfusion in the hippocampus following cessation of the stimulus. These effects were not observed using gamma stimulation and were still present even under conditions of pharmacologic NMDA antagonism. It is worth noting that arguably the most effective form of neuromodulation is still electroconvulsive therapy (ECT), which is performed under anesthesia and also results in increases to cerebral blood flow [55,56]. However, ECT is very non-specific, this lack of specificity may well contribute to its detrimental effect on memory [57]. Alternatively, transcranial magnetic stimulation (TMS) is most effective when applied focally to awake, alert patients [58,59]. Because DBS can combine a relatively high degree of modulation in deep structures with greater spatial and temporal specificity than ECT or TMS, it is plausible that DBS may have more benefits beyond that of ECT or TMS in treating disorders of cognitive function.

### Limitations and future directions

While the current study was performed in anesthetized animals, futures studies will investigate the effects of reduced NMDA function and MSN DBS in awake, behaving animals during memory-associated behavioral tasks (e.g., novel object recognition and Barnes Maze). The goal will be to determine if the observed ΔCBV within the septo-hippocampal network following theta-frequency MSN DBS is also associated with improved memory function, linking the present study with our previous study demonstrating improved memory following MSN DBS in MK-801 treated animals [27]. Another limitation of our study is that fUSI recordings performed using the conventional 1-dimensional linear ultrasound transducer array necessarily generates 2-dimensional pD vascular maps of the animals’ CBV. As a result, other regions that are connected with the MSN besides the hippocampus and mPFC (e.g., amygdala, habenula, raphe nucleus) were not accessible from the selected sagittal 2-dimensional image plane. Recent studies are tackling this challenge using whole-brain 3-dimensional fUSI with either moving linear arrays (similar to the array used in our study), matrix arrays or raw column arrays (RCAs) [60–62]. Future studies can use these probes to cover volumes rather slices of the mouse brain providing access to all areas of the septohippocampal network. Overall and regardless of these limitations, our findings demonstrate the feasibility of using fUSI to characterize network-specific neurovascular changes in disease models as well understand what parameters in neuromodulatory techniques most impact cerebral perfusion dynamics.

